# Modeling the role of immune cell conversion in the tumor-immune microenviroment

**DOI:** 10.1101/2023.03.22.533789

**Authors:** Alexander S. Moffett, Youyuan Deng, Herbert Levine

**Author notes:** Contributing authors.

## Abstract

Tumors develop in a complex physical, biochemical, and cellular milieu, referred to as the tumor microenvironment. Of special interest is the set of immune cells that reciprocally interact with the tumor, the tumor-immune microenvironment (TIME). The diversity of cell types and cell-cell interactions in the TIME has led researchers to apply concepts from ecology to describe the dynamics. However, while tumor cells are known to induce immune cells to switch from anti-tumor to pro-tumor phenotypes, this type of ecological interaction has been largely overlooked. To address this gap in cancer modeling, we develop a minimal, ecological model of the TIME with immune cell conversion, to highlight this important interaction and explore its consequences. A key finding is that immune conversion increases the range of parameters supporting a co-existence phase in which the immune system and the tumor reach a stalemate. Our results suggest that further investigation of the consequences of immune cell conversion, using detailed, data-driven models, will be critical for greater understanding of TIME dynamics.

## 1 Introduction

Cancer is a class of evolving diseases, similar in many ways to infectious diseases characterized by viral, bacterial, or eukaryotic parasite infection. Tumor cells can be thought of as cheaters in the cooperative multicellular state from which they are derived, although tumor cells often act collectively in a proto-multicellular manner [1, 2]. The key point is that cancer cells overcome controls ensuring intercellular cooperation and develop into a distinct population upon which evolutionary forces act, not unlike a population of pathogens within a host organism. While this evolutionary nature of cancer has long been acknowledged [3], recent work has rapidly developed our understanding of cancer population dynamics by applying concepts and theoretical tools from evolutionary biology and ecology [4–7]. Advances in lineage tracing [8] and genomics [9] technologies have allowed for unprecedented understanding of the complex eco-evolutionary processes underlying tumor growth and metastasis, and theoretical approaches drawing from evolutionary and ecological theory [10–13] have enabled predictions of eco-evolutionary phenomena in cancer and interpretation of experimental data. However, much is still unknown about cancer population dynamics, with many complexities still unexplored.

Recent work has developed the concept of the tumor microenvironment (TME), referring to the biochemical, cellular, and physical context in which a tumor exists and how this context impacts tumor behavior [14]. To further emphasize the specific role played by the immune system, one can focus on the tumor-immune microenvironment (TIME). The TIME concept emphasizes that tumors exist in an ecological context of immune and other host cells which influence tumor growth and progression in a complex manner [15]. While there are several clear differences between the ecological interactions of tumors with host cells and traditionally studied interactions in ecology, such as those between predator and prey animals [16], concepts from ecology have nonetheless proved useful in understanding tumor-immune interactions. These concepts hold promise for further untangling the complexities arising from nonlinear, multi-directional interactions between adapting (through varying combinations of mutations and phenotypic plasticity) populations of cells [17].

While the ability of the immmune system to suppress tumor proliferation and metastasis has rightfully received considerable attention [18], our understanding of how tumors attempt to shape the immune system into cancer-tolerant or even cancer-promoting states remains incomplete [19]. While tumors are well-known to affect the metabolic and biochemical state of the TIME [15, 20], tumor cells are also able to influence immune cell phenotypes [21], a process known as immune cell conversion. A prime example of immune cell conversion is the polarization of macrophages between M1 and M2 phenotypes [22, 23]. In the M1 phenotype, macrophages produce tumor-suppressing molecules, including nitric oxide and reactive oxygen species, while M2 macrophages produce pro-tumor factors and promote angiogenesis [24, 25]. Other examples of tumor-influenced immune cell conversion include T cell/regulatory T cell polarization [26] and NK cell/ILC1 cell polarization [27].

Despite strong evidence for the importance of immune cell conversion in the TIME, this phenomenon has been largely ignored in mathematical models of the TIME. While a number of models exist which include the polarization of immune cells in the TIME [28, 29], very few [30] have included the ability of tumor cells to bias this immune cell polarization. In order to explore the role of immune cell conversion in the TIME, we develop here a minimal mathematical model for tumor-immune interaction including ability of tumor cells to convert immune cells into a pro-tumor phenotype. Using modified Lotka-Volterra equations, we explore the effects of the rate of immune cell conversion, finding that conversion can be essential to the viability of a tumor population. Specifically, non-zero immune conversion rates can allow for tumor survival in the presence of non-trivial anti-tumor immunity. Our results highlight the need to further inspect the role of tumor-to-immune system feedback, especially in mathematical and computational models of cancer. Furthermore, greater understanding of the consequences of immune cell conversion may have an impact on the development of novel cancer immunotherapies.

## 2 Model

### 2.1 Guiding principles for tumor-immune modeling

Following the work of Wilson & Levy [31] and Arabameri *et al*. [32], we adopt a minimal description of the essential aspects of tumor-immune population dynamics by considering the densities of tumor, anti-tumor immune, and pro-tumor immune cells. This coarse-grained approach greatly simplifies our model and its analysis. This simplification is of course at the expense of the potential quantitative accuracy of a more detailed approach. Once we have established basic mechanisms, future efforts can extend our approach to include a larger number of cell types and We proceed by adapting the Lotka-Volterra framework [33] with multiplicative interaction terms. The resultant ODEs ignore both spatial aspects of tumor-immune interaction and complexities such as saturating interactions. We note that some TIMEs may be better represented by spatially homogeneous models than others. For example, tumor cells in leukemia are largely suspended in the bloodstream, and therefore the assumption that all cells interact with all other cells is a reasonable approximation. For solid tumors, our model assumes that is no barrier to immune infiltration [34]. Regardless of which biological scenarios our model is better suited for, we emphasize that our goal is to explore the potential consequences of immune conversion, not to make quantitative predictions about tumor-immune population dynamics.

### 2.2 A modified Lotka-Volterra model

We examine a generalized Lotka-Volterra model describing tumor cell (*T*), pro-tumor immune cell (*P*), and anti-tumor immune cell (*A*) population densities. Our basic innovation is the inclusion of a tumor-induced switching term from anti-tumor to protumor immune phenotypes. We abbreviate pro-tumor immune cells as PTI cells and anti-tumor immune cells as ATI cells. Our baseline model can be written as

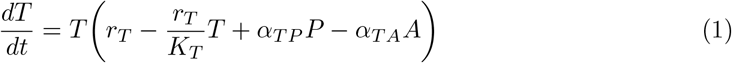

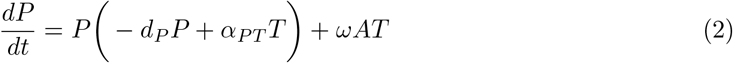

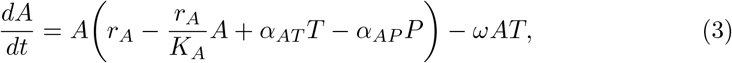

or in condensed vector notation

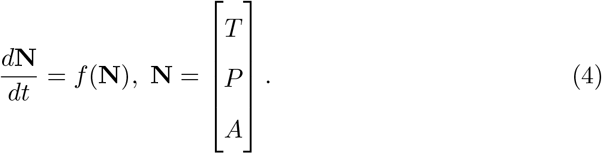

All parameters are non-negative, so that a negative sign in front of a parameter in Eqs. 1-3 indicates either inhibition of growth or contribution to death, while a positive sign indicates contribution to growth or inhibition of death. The parameters *r*_*T*_ and *r*_*A*_ describe the “intrinsic” growth rates of tumor and ATI cells each in the absence of other cell types, while *K*_*T*_ and *K*_*A*_ denote their carrying capacities. By calling *r*_*T*_ and *r*_*A*_ intrinsic growth rates, we mean that they are the growth rates in the absence of other interactions in the model. We assume that PTI cells mostly arise through tumor-induced conversion processes, so we take *r*_*P*_ to be zero, and denote their density-dependent growth inhibition and/or death rate as *d*_*P*_. The parameters *α*_*XY*_ represent the quadratic contributions of interactions between cell types to growth rates, where *α*_*XY*_ is the effect of *Y* on the net growth rate of *X*. Note that we assumed that contact with tumor cells induces growth of all types of immune cells.

Finally, as mentioned above, we include terms reflecting the ability of tumor cells to induce some immune cells that inhibit tumor growth, such as M1 macrophages, to switch phenotypes into functionally pro-tumor states, such as M2 macrophages. This is reflected in Eqs. 1-3 by a conversion parameter *ω* ≥ 0 controlling the rate at which tumor cells induce ATI cells to switch to PTI cells. We assume that all of these parameters are independent of time and cell concentrations.

### 2.3 Types of steady states

We refer to states of the system as feasible when the densities of all cell types are non-negative

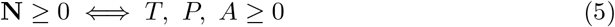

corresponding to physically meaningful states of the system. Eqs. 1-3 can support the following types of feasible steady states beyond the trivial steady state *T* = *P* = *A* = 0:

(**T**) Tumor-only: *T >* 0 and *P* = *A* = 0

(**A**) ATI-only: *A >* 0 and *T* = *A* = 0

(**TA**) Tumor-ATI coexistence: *T, A >* 0 and *P* = 0

(**TP**) Tumor-PTI coexistence: *T, P >* 0 and *A* = 0

(**TPA**) Tumor-PTI-ATI coexistence: *T, P, A >* 0.

Tumor-only (**T**) and ATI-only (**A**) steady states always exist and are always feasible in the allowed parameter space, while the simultaneous feasibility and existence of steady states **TA, TP**, and **TPA** depend on the choice of parameters. As we will see, there can be at most one steady state corresponding to types **T, A, TA**, and **TP**, while there can be more than one tumor-PTI-ATI coexistence (**TPA**) steady state.

We denote the set of steady state solutions corresponding to each type as 𝒮_*X*_, where *X* in the subscript indicates the cell types that are positive in the steady state. For example, for the tumor-only steady states, we have

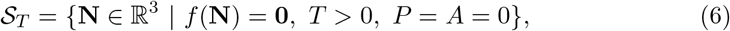

while for tumor-PTI-ATI coexistence steady states we have

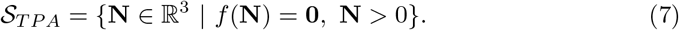

Because there can be at most one feasible steady state for each of the types (I)-(IV), we can unambiguously refer to *the* steady state meeting the respective criteria of these types, writing these as

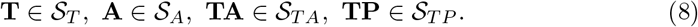

We can refer to steady states with tumor-PTI-ATI coexistence similarly

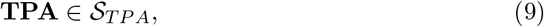

but we must often take care to specify which solution in S_*T P A*_ we are referring to.

### 2.4 Linear stability analysis

In order to assess the linear stability of steady states, we use the Jacobian matrix

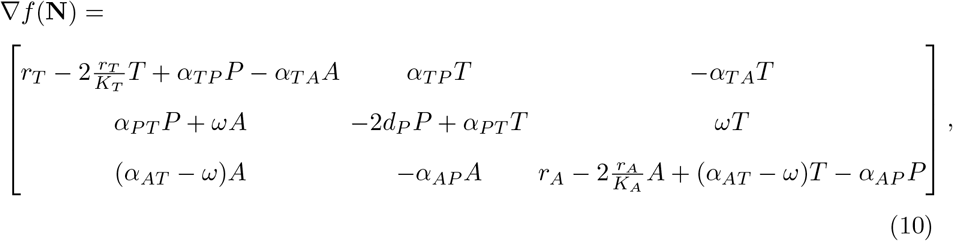

obtained from taking partial derivatives of the right-hand side of Eqs. 1-3 with respect to *T, P*, and *A*, and check whether the real parts of its eigenvalues are all non-positive. We call steady states stable when this is the case.

### 2.5 Computational methods

We performed all numerical ODE integration and root finding using SciPy 1.7 [35]. We used NumPy 1.21 [36] in most calculations. For creating plots, we used Matplotlib 3.5 [37] within a Jupyter notebook [38]. We used the scikit-learn 1.0 [39] support vector machine implementation with a degree 3 polynomial kernel for visualizing the interfaces between basins of attraction in Fig. 4.

## 3 Results

### 3.1 Characterizing the steady states through feedback

As described above, we can classify the steady states of our model by the cell types that have non-zero density. For all classes of steady states, except for the case of tumor-PTIATI coexistence (**TPA**) with *ω >* 0, we can write the steady state densities in simple terms of the model parameters (Table 2). We can further simplify these expressions by gathering parameters into terms describing effective growth rates, written as Ω_*X→Y*_, meaning the net growth rate of *X* when “invading” a steady state population of *Y*, and total negative feedback, written as Γ_*X→Y* ;*S*_, meaning the total negative feedback of *X* on *Y* at the specified steady state *S*. For example, the effective growth rate of tumor cells in a population of ATI cells is

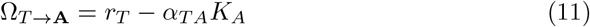

We note that by “invasion” we refer to the concept of invasive populations in ecological settings, not implying anything that is occurring spatially. Similarly, the total negative feedback of tumor cells on tumor cells in the tumor-PTI-ATI coexistence state (**TPA**) with *ω* = 0 is

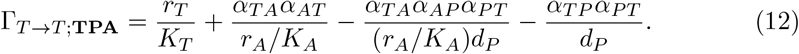

Note that the3 feeback depend directly on the state *S*, because zero density of one or more cell types will remove feedback “channels”, limiting the number of ways that feedback can be felt by a cell type. To illustrate this, we can compare the total feedback of tumor cells on tumor cells in the tumor-PTI-ATI coexistence state (Eq. 12) with that for the tumor-PTI coexistence state (**TP**)

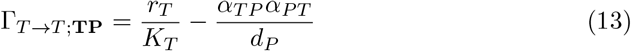

which lacks all interaction parameters with ATI cells (see Fig. 1).

**Fig. 1.**
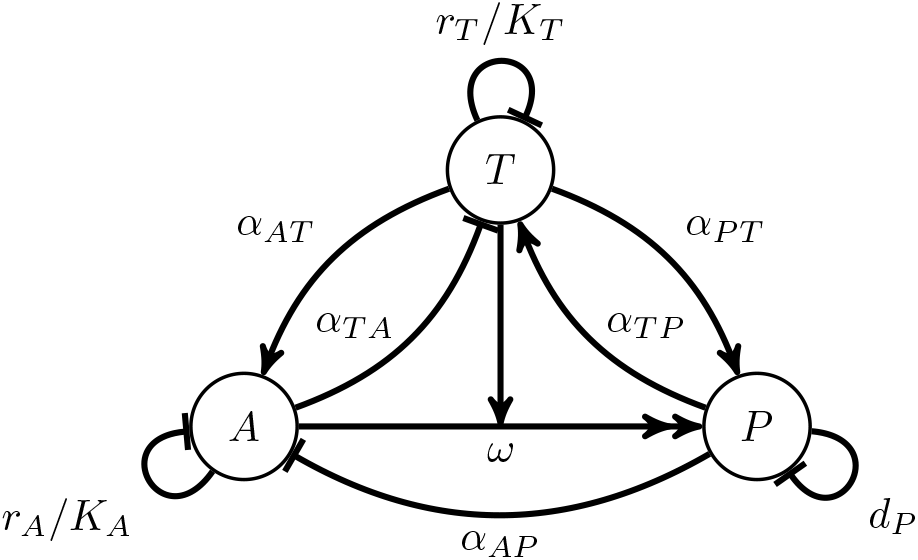
Schematic diagram of interactions in the model. Note that the exponential growth terms *r*_*T*_ *T* and *r*_*A*_*A* are not depicted. The meanings of each type of line between *T, P*, and *A* are as follows. Single pointed arrow: positive (“activating”) interaction; Line with straight, perpendicular end: negative (“inhibiting”) interaction; Double pointed arrow with perpendicular, bisecting single pointed arrow: conversion interaction, here conversion of A to P is “catalyzed” by T.

The tumor-only (**T**) and tumor-ATI coexistence (**TA**) steady states are always unstable (Table 2), and are therefore of no interest for our purposes. This leaves the ATI-only (**A**), tumor-PTI coexistence (**TP**), and tumor-PTI-ATI coexistence (**TPA**) states as the focus of our analysis. These can be thought of respectively as immune “wins”, tumor “wins”, and tumor-immune “draw”. To narrow the parameter space of interest, we can see that **TP** is feasible only when

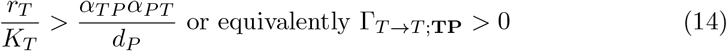

is satisfied. This can be interpreted to mean that the direct negative feedback of tumor cells on themselves must be greater than the positive feedback of tumor cells on themselves through PTI cells, where the feedback signs are clear from Eqs. 1 & 2 (tumors are self-limiting through *r*_*T*_ */K*_*T*_ while tumor cells increase PTI growth rate and PTI cells increase tumor growth rate). If this condition is violated, unbounded tumor growth is possible, contradicting the biophysical realities of cancer. Thus, we can reasonably focus on the parameter space where Eq. 14 is met, as we do throughout the remainder of this work.

### 3.2 Stability of steady states

The stability conditions we have derived for several of the steady states have several interesting implications. The **A** state is only stable when the tumor invasion growth rate (Ω_*T →***A**_) is non-positive (Table 2). When *ω* = 0, a necessary condition for the **TPA** state to be stable is that Γ_*T →T* ;**TPA**_ *>* 0 (Appendix A). This means that the only way for there to be a feasible, stable tumor-PTI-ATI steady state when *ω* = 0 is for Ω_*T →***A**_ to be positive (Table 2). Thus, when *ω* = 0 the **A** and **TPA** steady states cannot both be feasible and stable. There can, however be bistability for *ω* = 0 between the **A** and **TP** states.

The stability of the **TP** state can switch when *ω* is increased, provided that the right-hand side of the stability condition

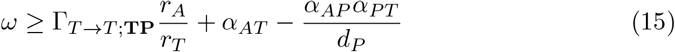

is positive. We will see in the next section that this stability switch corresponds to a transcritical bifurcation where the **TPA** state collides with the **TP** state. The **TP** state can only be stable for *ω* = 0 when the right-hand side of Eq. 15 is non-positive (see Appendix A).While we can exactly solve for the **TPA** state and the eigenvalues of the corresponding Jacobian for any *ω* value, the complexity of the resulting expressions prevent clear interpretation. See Appendix A for a more detailed account of solving for **TPA**.

### 3.3 Viability of tumor cell populations is dependent on immune cell conversion

Given the stability conditions discussed in the previous section, there are several possible scenarios exhibited by the tumor-immune ecosystem when immune cells cannot be converted (*ω* = 0). There can be a single, stable, steady state, either **A** or **TPA**, or there can be bistability between **A** and **TP**. This bistability can only occur when ATI cells cannot invade the **TP** state, an exceptionally hostile environment for the normally functioning immune system. Such a situation seems unlikely except for well-established tumors in decidedly pro-tumor TIMEs.

Of more interest is the behavior found when including finite *ω* effects. The top row of Fig. 2 shows the behavior of the steady states as *ω* varies when Ω_*T →***A**_ *>* 0, where an increasing killing rate of tumor cells by ATI cells (*α*_*T A*_) decreases the tumor density in **TPA**. When Ω_*T →***A**_ = 0, as shown in the second row of Fig. 2 (lightest green curve in the subplot), the stable **TPA** solution disappears altogether at *ω* = 0, while the **A** state becomes stable. When there is no stable state with a non-zero tumor cell density at *ω* = 0 (Ω_*T →***A**_ *<* 0), a saddle node bifurcation occurs at a positive *ω* (Fig. 2). Above this *ω* value, there are two **TPA** steady states, one stable and one unstable. As *ω* increases, eventually the stable **TPA** solution collides with the **TP** state, exchanging stability in a transcritical bifurcation. For larger *ω* values, past the transcritical bifurcation, there is bistability between the **A** and **TP** states.

**Fig. 2.**
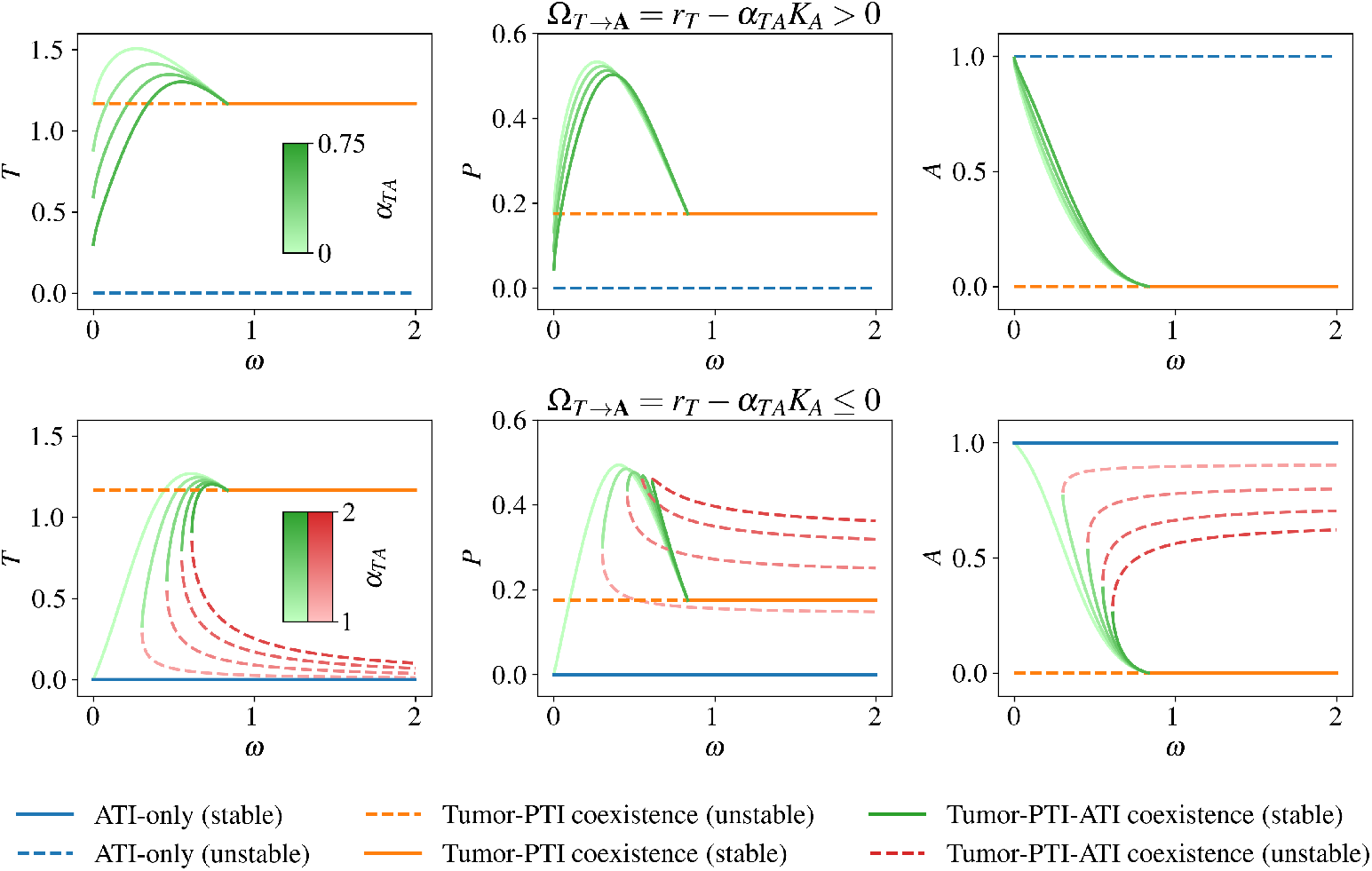
Bifurcation diagrams over *ω* for different ATI tumor-killing rates (*α*_*T A*_). When tumor cells can invade an ATI-only state (Ω_*T →***A**_ *>* 0, first row), there is no stable steady state with *T* = 0. The **TPA** state has increasing values of *T* and *P* as *ω* increases, until it collides with the **TP** state in a transcritical bifurcation, swapping stabilities. On the other hand, when tumor cells cannot invade an ATI-only state (Ω_*T →***A**_ *≤* 0, second row), there is always a stable cancer-free state **A**. If Ω_*T →***A**_ = 0 (*α*_*T A*_ = 1 in the second row, with the **TPA** steady state shown in the lightest shade of green), then at *ω* = 0 the only stable steady state is **A**, with the **TPA** steady state appearing for *ω >* 0. In this case, there is a saddle node bifurcation at a value of *ω* below which there is no steady state with positive tumor density. Above this value of *ω*, there is bistability between the cancer-free and cancer states. The first column shows the steady state densities of tumor cells, while the second column shows the density of PTI cells, and the third column shows the density of ATI cells. Only feasible steady states are shown. Note that each individual subplot is a projection onto to *T, P*, or *A* densities from the full three-dimensional state space, so that intersections of solutions occur only when curves in all three projections intersect.

The saddle node bifurcation occurs at larger *ω* values as *α*_*T A*_ increases, meaning that in order to maintain a positive steady state population density, tumor cells must convert ATI cells more rapidly when ATI cells kill tumor cells more efficiently. When *α*_*T A*_ is large enough, the stable branch of the **TPA** state disappears and a pitchfork bifurcation occurs at 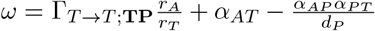 (Fig. 3). For even larger values, the *ω* value at which the saddle node bifurcation occurs decreases, but the stable **TPA** steady state is no longer feasible.

**Fig. 3.**
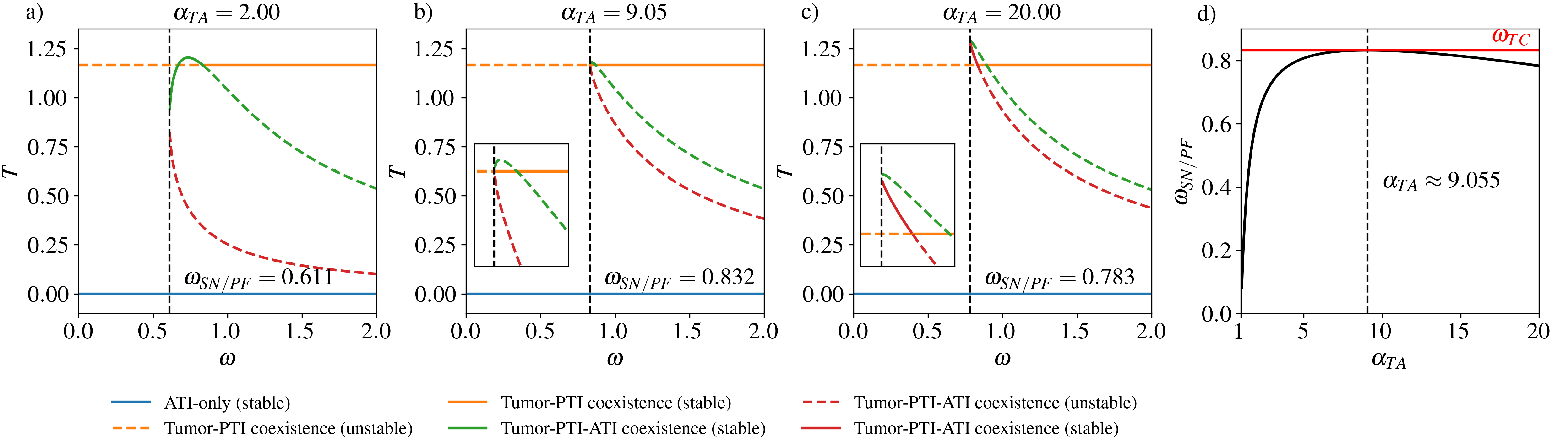
Values of *ω* for saddle node and pitchfork bifurcations. a)-c) Bifurcation diagrams showing the steady state tumor cell density as *ω* is changed, as in Fig. 2, for three representative *α*_*T A*_ values. a) For *α*_*T A*_ = 2, we see the same behavior as in the last row of Fig. 2. b) At *α*_*T A*_ *≈* 9.05, there is a pitchfork bifurcation rather than a saddle node bifurcation. The inset subfigure shows a zoomed-in view of the pitchfork bifurcation. c) For the large value of *α*_*T A*_ = 20, there is again a saddle node bifurcation, which occurs at a value of *ω* which now decreases as *α*_*T A*_ is further increased. The stable part of the lower red branch of the tumor-PTI-ATI solution (before colliding with the tumor-PTI solution in a transcritical bifurcation) is not feasible. The inset subfigure shows a zoomed-in view of the saddle node and transcritical bifurcations. d) The value of *ω* at which the saddle node bifurcation (or pitchfork bifurcation in the special case of *α*_*T A*_ *≈* 9.055) occurs, labeled as *ω*_*SN/PF*_, as a function of *α*_*T A*_. The *ω* value where the tumor-PTI-ATI solution and the tumor-PTI solution collide in a transcritical bifurcation (*ω*_*T C*_ *≈* 0.832) is shown as a red horizontal line.

There are several interesting biological implications of these results. First, we would expect to see a minimal immune cell conversion rate for tumor viability, below which the tumor cannot be sustained. We would not expect to observe this phenomenon in all cases, as we see viable tumor cell populations at *ω* = 0 for tumor-friendly parameters. Rather, we would expect to find minimal immune conversion rates for tumor viability in the early stages of tumor development. This is a testable prediction of our model. If the production rate or degradation rate of the biochemical messengers mediating immune cell conversion, such as TGF*β* [21] can be experimentally manipulated, it should be possible, albeit perhaps technically difficult, to examine the effects of *ω* on tumor viability *in vitro*.

Second, our model suggests that a “stalemate” state (**TPA**) can exist at intermediate immune conversion rates. The steady state with coexistence between tumor cells, PTI cells, and ATI cells represents a stalemate between pro-tumor and anti-tumor cell types, consistent with previously hypothesized “equilibrium” tumor states [40]. In the context of metastasis, this possibility is sometimes referred to as tissue dormancy. We predict that, intuitively, the possible sizes of “equilibrium” tumor populations are limited by the ability of ATI cells to kill tumor cells (*α*_*T A*_, see Fig. 2).

Third, we find that there can be an intermediate immune cell conversion rate at which the tumor population density is maximal. This is due to the fact that the steady state ATI cell “reservoir” for producing PTI cells shrinks as *ω* increases, so that in some parameter sets there is an optimal *ω*_max_ such that for larger *ω > ω*_max_, the tumor cell density is less than at *ω*_max_. This phenomenon occurs as the stable **TPA** solution approaches the **TP** solution, in which the anti-tumor immune system is nonexistent in the local TIME. While it is unclear how well a complete (local) lack of ATI cells reflects a later stage TIME, the possibility of an intermediate immune cell conversion rate that maximizes tumor cell density has not, to our knowledge, been discussed previously.

Finally, our model predicts that above a threshold immune cell conversion rate, large tumor killing rates by ATI cells (*α*_*T A*_) are not sufficient to eradicate stable states with positive tumor density (Fig. 3). In this way, immune cell conversion can protect tumor viability against even an extraordinarily effective immune system.

### 3.4 Immune cell conversion promotes tumor survival at small growth rate

As discussed in the previous section, our model indicates that when Ω_*T →***A**_ = *r*_*T*_ −*α*_*T A*_*K*_*A*_ ≤ 0, there cannot be a stable **TPA** state when *ω* = 0. While we have focused on the effects of *α*_*T A*_ in Fig. 2, clearly reducing *r*_*T*_ (reflecting a decreased tumor growth rate) or increasing *K*_*A*_ (reflecting a larger ATI cell carrying capacity) can yield similar results. When the intrinsic tumor growth rate (*r*_*T*_), the rate of ATI cells killing tumor cells (*α*_*T A*_), and the ATI cell carrying capacity (*K*_*A*_) yield a negative Ω_*T →***A**_, we observe bistability between **A** (cancer-free) and either **TPA** or **TP** (cancer), provided that *ω* is large enough. This leads to basins of attraction for the cancer-free and cancer states, divided by an *ω*-dependent two-dimensional surface (Fig. 4).

**Fig. 4.**
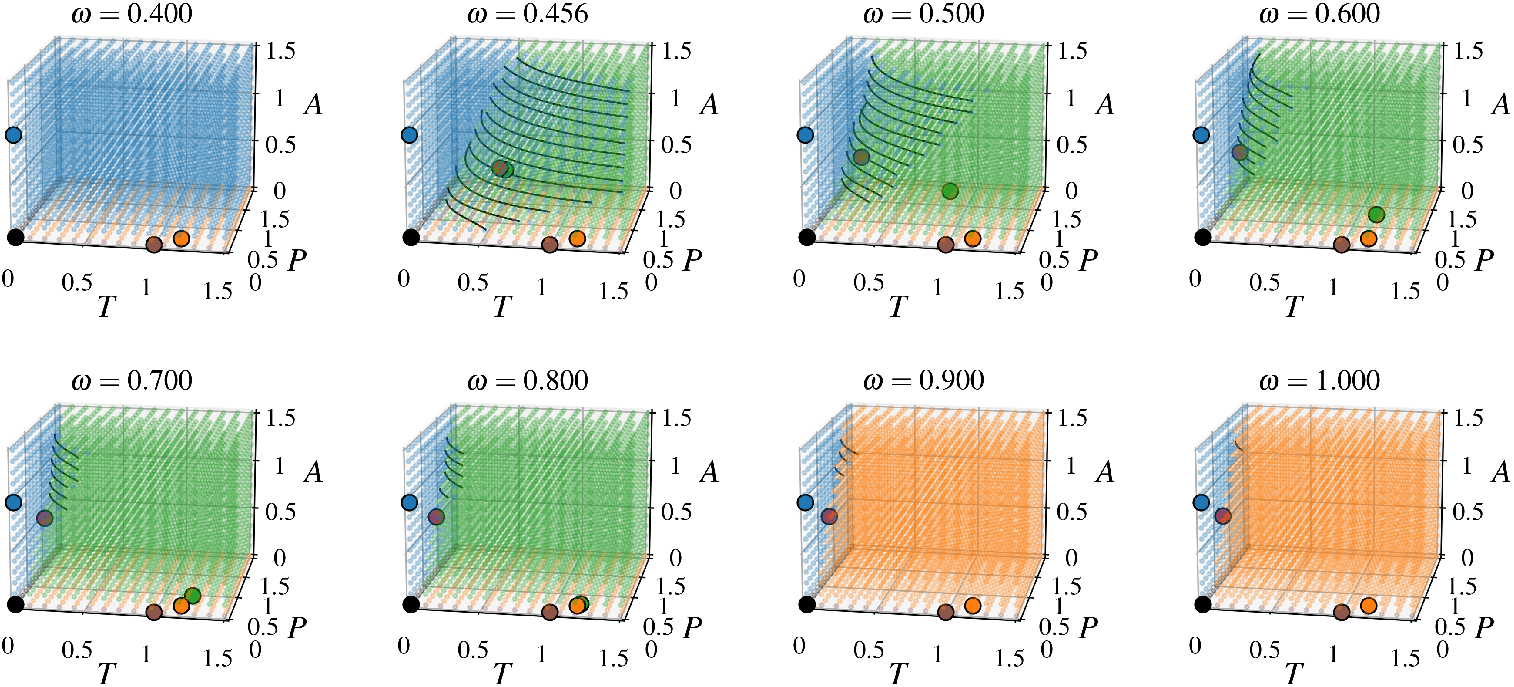
The basins of attraction with increasing *ω*. As *ω* increases, the region of initial states that end up at the cancer-free state diminishes rapidly, while the basins of attraction for states with positive tumor density increase in volume. Each subplot depicts a three-dimensional grid of points representing initial values of *T, P*, and *A*. The larger dots with black edges indicate the locations of steady states, with colors matching those in Fig. 2 (note that the *T* = *P* = *A* = 0 and **T** steady states, shown in black and brown respectively, are not shown in Fig. 2). The points in the three-dimensional grid are colored according to the steady state that they asymptotically approach in numerical integration of Eqs. 1-3. The black curves are estimates of the dividing surface contours separating basins of attraction, found using support vector machines with degree 3 polynomial kernels as implemented in scikit-learn 1.0 [39].

This bistability between a state with zero tumor density and a state with positive tumor density is an example of an Allee effect, a term commonly used in ecology [5]. With an initially low tumor density, the system will evolve towards the cancer-free state, while with sufficiently large tumor and ATI cell density the system will evolve towards the cancer state. As the ATI-to-PTI conversion rate increases, the region of state space where the system will evolve towards the cancer free state shrinks (Fig. 4). This means that if the system is initially in the cancer-free state, the amount of tumor density that must be introduced for the system to reach the cancer state decreases as *ω* increases.

Without any ATI-to-PTI conversion (*ω* = 0), there cannot be bistability between the **A** and **TPA** states. However, there can still be bistability between **A** and **TP** when

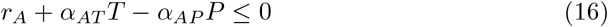

where *T* and *P* are the steady state densities in the **TP** state (see Eq. 15). This means that the growth rate of a small ATI density introduced to the **TP** state must be negative (Eq. 2), consistent with developed TIME state that is uninhabitable for ATI cells. Thus, while an ATI-to-PTI conversion term is not strictly necessary for bistability, it allows for bistability in a broader range of parameter sets with biological relevance.

Finally, we note that this baseline model does not exhibit tristability. That is, the existence of a co-existence state means that for some range of parameters further growth of the tumor is precluded by an increasing immune response. There is no mechanism whereby even a large increase in tumor size could overcome this linear response, hypothetically giving rise to **A, TPA, TP** tristability. We cannot exclude the possibility that a more complete model might exhibit such a parameter region.

### 3.5 Alternate modeling choices yield similar results

While we have sought to analyze a minimal model of tumor-immune interaction, there are several alternate modeling choices we could have made, depending on assumptions about the behavior of the immune system. In order to test the effects of changing our assumptions, we consider three modifications to our original model in Eqs. 1-3

1. Linearity of PTI direct self feedback: −*d*_*P*_ *P*^2^ or −*d*_*P*_ *P* in Eq. 2
2. Direct positive feedback from tumor cells to PTI cells: *α*_*P T*_ *>* 0 or *α*_*P T*_ = 0 in Eq. 2
3. ATI proliferation and non-linear direct self feedback or constant recruitment with linear direct self feedback: 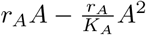 or 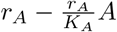 in Eq. 3.

Altogether, there are eight total models spanned by all the choices listed above, leading to seven alternate models to our original set of equations in Eqs. 1-3. With the same parameters examined for the original model (Table 1), we find that the bifurcation behavior remains qualitatively similar for all of the eight models (Fig. 5). However, there are several noticeable differences with the alternate models. One of these differences is that for each model except for the original, there is no stable, feasible **TP** state for any *ω* value. Additionally, when ATI dynamics are altered so that ATI cells are recruited in an *A*-independent manner and die at a constant rate, there is no **T** state, so that the tumor density of the **TPA** state, when it exists, is an increasing function of *ω*. For a more detailed analysis of the alternate models, see Appendices B & C.

**Table 1.** Parameter values used in figures.

| Parameter | Fig. 2 | Fig. 3 | Fig. 4 | Figs. 5-7 |
| --- | --- | --- | --- | --- |
| $r_T$ | 1 | 1 | 1 | 1 |
| $K_T$ | 1 | 1 | 1 | 1 |
| $d_P$ | 1 | 1 | 1 | 1 |
| $r_A$ | 1 | 1 | 1 | 1 |
| $K_A$ | 1 | 1 | 1 | 1 |
| $\alpha_{TP}$ | 0.95 | 0.95 | 0.95 | 0.95 |
| $\alpha_{TA}$ | Varying in $[0, 2]$ | Varying in $[1, 20]$ | 1.5 | 1.5 |
| $\alpha_{PT}$ | 0.15 | 0.15 | 0.15 | Varying in $\{0, 0.15\}$ |
| $\alpha_{AT}$ | 0.05 | 0.05 | 0.05 | 0.05 |
| $\alpha_{AP}$ | 0.5 | 0.5 | 0.5 | 0.5 |
| $\omega$ | Varying in $[0, 2]$ | Varying in $[0, 2]$ | Varying in $[0, 1]$ | Varying in $[0, 5]$ |

**Table 2.**
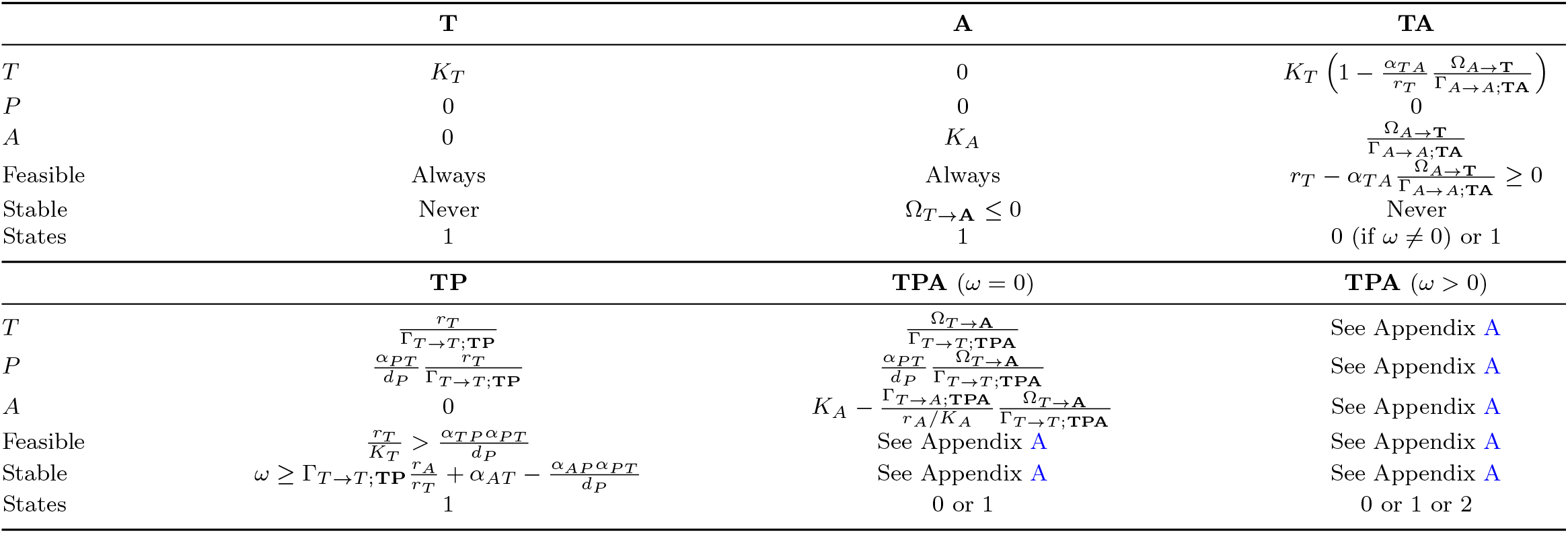
Summary of steady states. See Appendix A for detail.

**Fig. 5.**
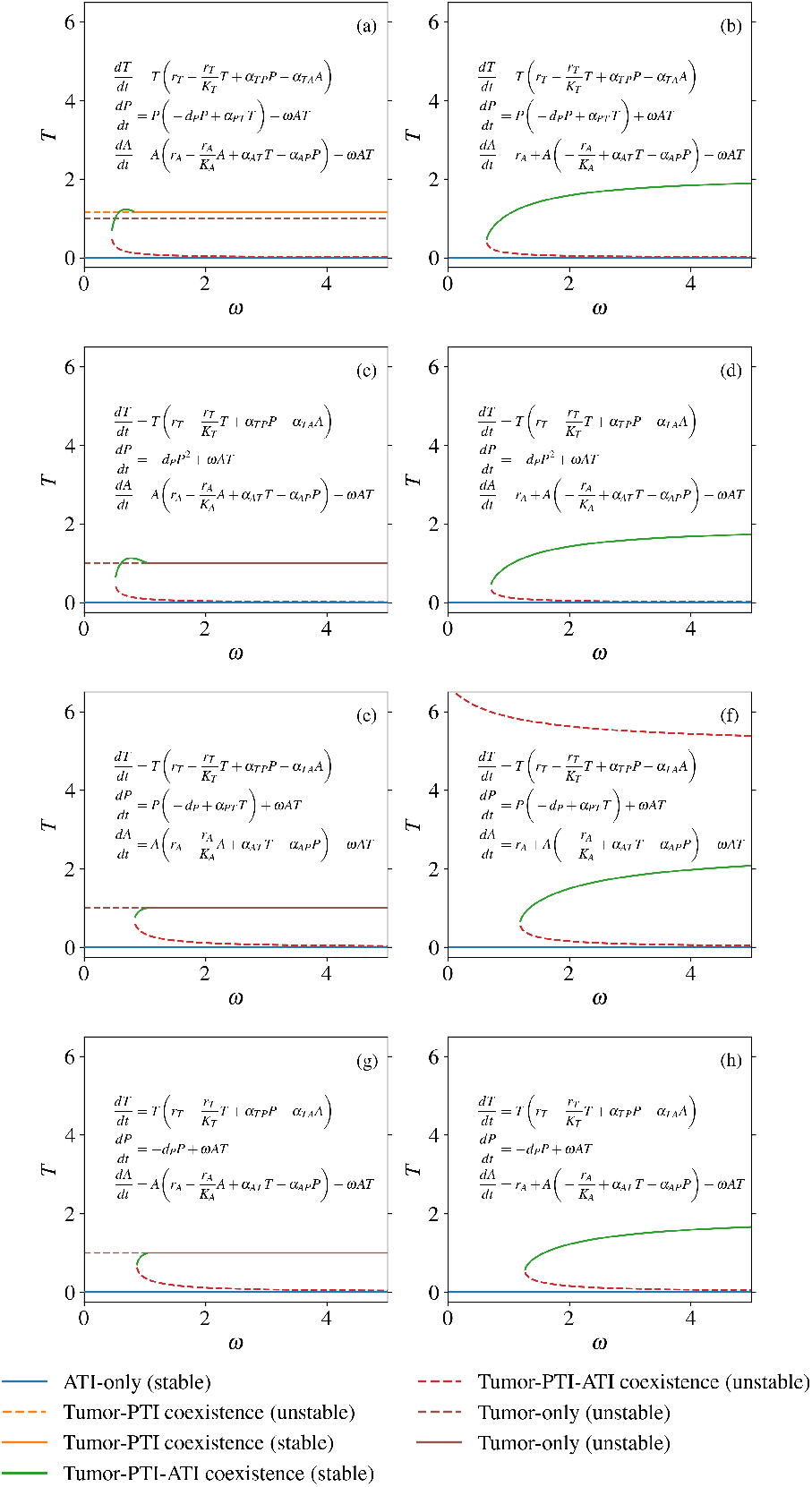
Bifurcation diagrams showing the steady state density of tumor cells from all eight choices of model. (a) The model from Eqs. 1-3, analyzed in the above sections. (b) The model in (a) except with an ATI source term (immune cell recruitment) and a linear ATI death term. (c) The model in (a) without a PTI growth term dependent on tumor cells (equivalent to *α*_*P T*_ = 0). (d) The model in (c) with an ATI source term and a linear ATI death term. (e) The model in (a) with a linear PTI death term. (f) The model in (d) with an ATI source term and a linear ATI death term. (g) The The model in (a) without a PTI growth term dependent on tumor cells and with a linear PTI death term. (h) The model in (g) with an ATI source term and a linear ATI death term. With a linear intrinsic ATI birth term and non-linear ATI death rate, in subplots (a), (c), (e), and (g), the stable tumor-PTI-ATI steady state collides either with the tumor-PTI coexistence state in (a) or with the tumor-only state in (c), (e), and (g). On the other hand, with an ATI source term and a linear ATI death term, in subplots (b), (d), (f), and (h), the stable tumor-PTI-ATI steady state does not collide with another steady state but saturates as *ω* increases.

When PTI cells have a non-linear self-limitation term, the saddle node bifurcation leading to bistability occurs at a lower *ω* value (Fig. 5). This difference is likely due to the fact that the value of *P* in the **TPA** state for the parameters we used is less than one, leading to less self-limitation than would occur with a linear self-limitation term. Thus, the saddle node bifurcation will likely occur at a larger *ω* value for these models than for those with linear PTI self-limitation terms with the same parameters.

## 4 Conclusions

We have analyzed the effects of tumor-induced immune cell conversion in a simple model of the TIME, finding that an immune cell conversion term allows for bistability between a cancer-free state and a state with a non-zero tumor cell density. Our results suggest an important role for immune cell conversion in the early stages of tumor growth, before the TIME has been shaped into a pro-tumor state, which in the context of our model is characterized by parameters for which there is no stable steady state with positive tumor density for *ω* = 0. For a large enough immune cell conversion rate, we find that a “stalemate” or “equilibrium” stable steady state can exist. In this stalemate state, tumor cells, PTI cells, and ATI cells can coexist, maintained by a balance of pro-tumor and anti-tumor factors. Eventual escape from equilibrium tumor states, leading to tumor growth not limited by the anti-tumor immune system, is not directly captured by our model; perhaps including a direct competition for metabolic resources might allow a large enough tumor to completely suppress immunity. Instead, escape from a coexistence state can be caused by changes in model parameters so that a shift from a coexistence state to a stable tumor and PTI cell steady state occurs.

By assuming a quadratic form for ATI-induced tumor cell death (−*α*_*T A*_*TA* in Eq. 1), we ignore the possibility of tumor cell population size-dependent or tumor growth rate-dependent immunosurveillance [41]. In the case where small tumor cell populations are not detected and/or targeted by the immune system, it is conceivable that a large enough tumor cell population can grow before ATI-induced tumor cell death becomes appreciable, allowing tumors to bypass the basin of attraction for **A** states when there is bistability with steady states characterized by a non-zero tumor cell density. In this case, the tumor cell population density at which the immune system begins killing tumor cells can be considered the initial state of a tumor in our model, and larger ATI-to-PTI conversion rates will place this initial state closer to the basin of attraction of cancer states. At the same time, our model suggests that an immune system able to successfully reduce a large tumor cell density may be able to bring tumor cell density to a threshold value, below which tumor clearance is nearly inevitable.

Previous work has suggested that tumor cell populations may subject to an Allee effect [5, 42–44]. Allee effects can be classified as weak, where below a threshold population size the growth rate is non-negative but small, or strong, where below a threshold the growth rate is negative, driving the mean population size to zero. One suggested mechanism for an Allee effect in tumor population dynamics is cell-density dependence of “go or grow” phenotype switching [42]. Our ecological model of the TIME suggests that tumor-immune interactions, namely immune cell conversion, may also contribute to an Allee effect. Clearly, if a threshold initial population of tumor cells is necessary for the tumor to persist (on average), there is a significant barrier to tumor viability.

As the ability of tumor cells to convert ATI cells to PTI cells increases, the threshold population size decreases, lowering the barrier for tumor viability as a function of initial population size.

Further work is needed to uncover the role of tumor-induced immune cell conversion on cancer dynamics. From the theoretical side, more realistic treatment of the immune system, together with consideration for the effects of spatial organization on tumor growth and immune interaction could be incorporated into modeling efforts. In addition, phenotypic switching of tumor cells should be taken into account, especially since processes such as EMT might alter a cell’s sensitivity to immune interdiction [45]. Ideally, a quantitatively predictive model could be developed to allow for direct comparison with experiments, in order to test our theory-generated hypotheses concerning the role of immune cell conversion in tumor dynamics. Further, such a model could inform immunotherapy strategies targeting immune cell conversion, including attempts to promote M1 tumor-associated macrophage phenotypes over M2 phenotypes [30, 46–48] and the targeting of regulatory T-cells [49].

## Acknowledgments

This work was supported in part by the Center for Theoretical Biological Physics, NSF PHY-2019745.

## Data availability

Code needed to produce all results and figures from this article is available at: https://github.com/amoffett/tumor-immune_ecology_2023.

## Appendix A Analysis of steady states

### A.1 Tumor-only (T)

The tumor-only steady state (**T**) is

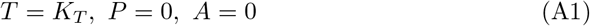

and is always feasible in the allowed parameter space. The eigenvalues of the Jacobian for **T** are

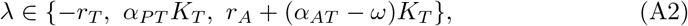

so that the tumor-only steady state is always unstable because *α*_*P T*_ *K*_*T*_ *>* 0.

### A.2 ATI-only (A)

The ATI-only steady state (**A**) is

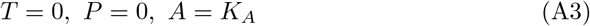

with eigenvalues of the Jacobian

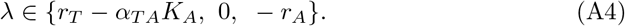

Then, the ATI-only steady state is stable only when

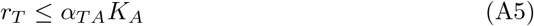

is satisfied. This means that for the cancer-free ATI-only state to be stable, the ability of ATI cells to reduce the growth rate of tumor cells at the maximal ATI population size (in the absence of other cell types) must outpace the intrinsic growth rate of tumor cells.

We can rewrite the condition in Eq. A5 by defining the invasibility of a healthy system in the ATI-only state by tumor cells as

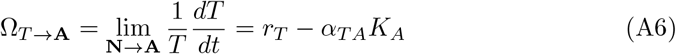

so that when Ω_*T →***A**_ *>* 0 tumor cells can invade and when Ω_*T →***A**_ ≤ 0 they cannot. We can restate the stability condition in Eq. A5 in terms of tumor invasibility as

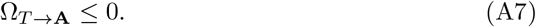

### A.3 Tumor-ATI coexistence (TA)

The tumor-ATI steady state (**TA**) can only exist when *ω* = 0, because otherwise the combination of tumor cells and ATI cells would produce PTI cells, adding PTI cell density. The steady state solution is

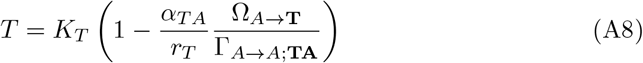

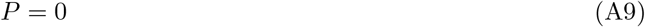

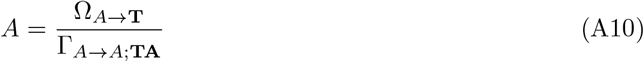

with the invasibility of the tumor-only state by ATI cells and the total negative feedback of ATI cells on themselves in the tumor-ATI coexistence steady state defined as

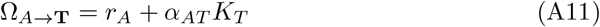

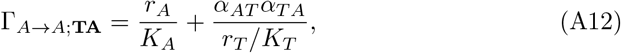

respectively. Note that both Ω_*A→***T**_ and Γ_*A→A*;**TA**_ are non-negative, so that the tumor-ATI coexistence steady state is feasible when

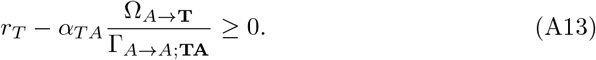

This condition can be interpreted to mean that the tumor-ATI coexistence steady state is feasible when the growth of the tumor outpaces tumor death due to ATI cells. The tumor-ATI coexistence state is unstable, as when a small density of PTI (*δP >* 0) is added to the system, the net growth rate of PTI cells (plugging *δP* into Eq. 2 with *ω* = 0)

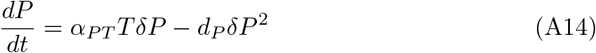

will be positive. It is clear that when

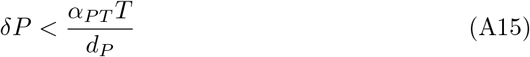

is satisfied Eq. A14 is positive, providing an explicit definition for a “small density” in this case.

### A.4 Tumor-PTI coexistence (TP)

The tumor-PTI coexistence steady state (**TP**) is

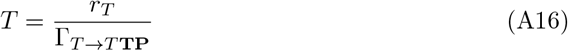

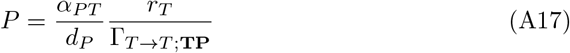

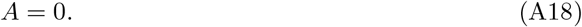

where the total negative feedback of tumor cells on themselves in this steady state is written as

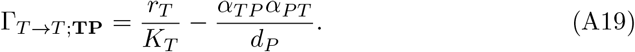

In order for tumor-PTI coexistence to be feasible, the negative feedback of tumor cells on themselves must be positive,

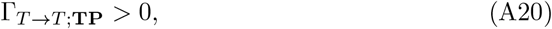

meaning that the direct, negative feedback of tumor cells on themselves must exceed, in magnitude, the positive feedback of tumor cells on themselves through PTI cells.

Unlike for previously discussed steady states, the stability of the tumor-PTI coexistence steady state is dependent on the ATI-to-PTI conversion rate *ω*. The eigenvalues of the Jacobian are

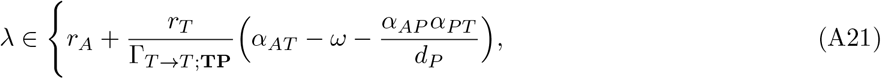

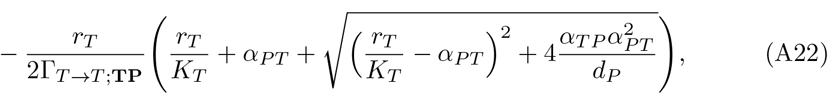

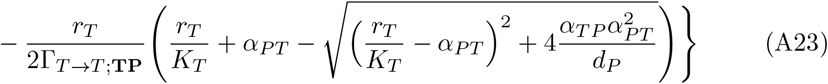

As long as

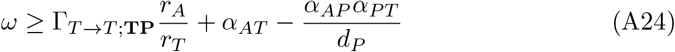

holds then the first eigenvalue (Eq. A21) will be non-positive. The second eigenvalue (Eq. A22) will always be non-positive, while the third eigenvalue (Eq. A23) be nonpositive only when the condition

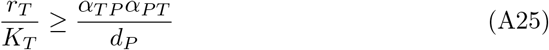

is met. However, this condition is identical to that required for feasibility of the tumor-PTI coexistence state (Eq. A20), except that feasibility requires strict inequality. Thus, when the tumor-PTI coexistence state is feasible, it is stable if and only if the condition in Eq. A24 is met. When the right-hand side of Eq. A24 is non-positive, the tumor-PTI coexistence steady state is stable for all *ω* ≥ 0, provided that Eq. A25 is also satisfied. We can write the condition of right-hand side of Eq. A24 being non-positive as

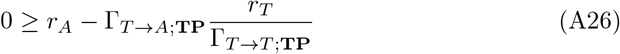

with

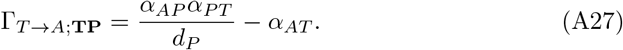

This corresponds to the invasion growth rate of ATI cells in the **TP** steady state, so that if ATI cells cannot invade then **TP** is stable for all *ω* ≥ 0.

### A.5 Tumor-PTI-ATI coexistence (TPA) without PTI to ATI conversion (*ω* = 0)

When *ω* = 0, the tumor-PTI-ATI (**TPA**) coexistence steady state is

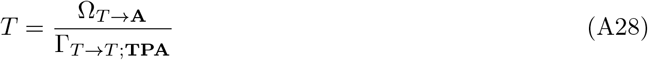

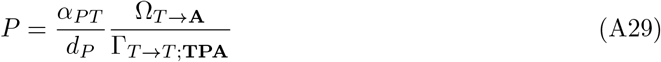

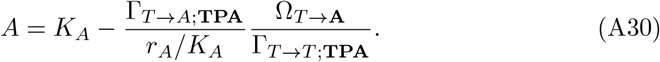

where the total negative feedback with zero ATI to PTI conversion from tumor cells on themselves and on ATI cells, respectively, is

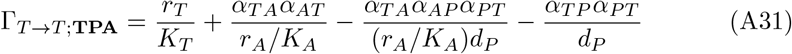

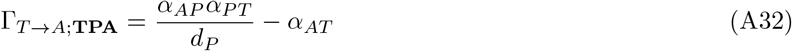

and Ω_*T →***A**_ is defined as in Eq. A6. Note that −*α*_*P T*_ can similarly be defined as the total negative tumor to PTI feedback.

Clearly, a necessary condition for the tumor-PTI-ATI coexistence steady state to be feasible is that the signs of Ω_*T →***A**_ and Γ_*T →T* ;**TPA**_ must be the same. If we perturb the steady state when Γ_*T →T* ;**TPA**_ *<* 0, however, we can see that it is unstable. For a small, positive perturbation of the tumor cell density (*δT >* 0), we define the overall perturbation

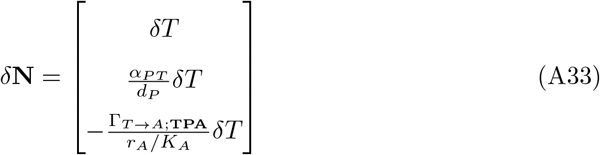

and then plug this into Eq. 1 for the change in tumor cell density, with *T, P*, and *A* taking on the values in Eqs. A28-A30, to find

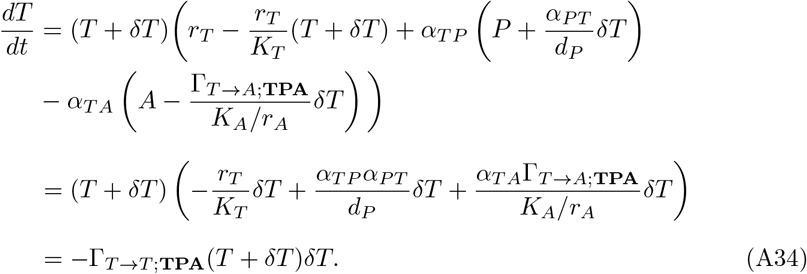

If Γ_*T →T* ;**TPA**_ *<* 0, as we have assumed, then the tumor cell density will increase, failing to restore the system to the steady state. Checking for *P*

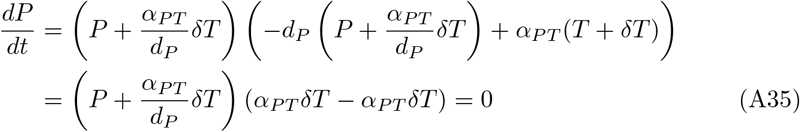

and *A*

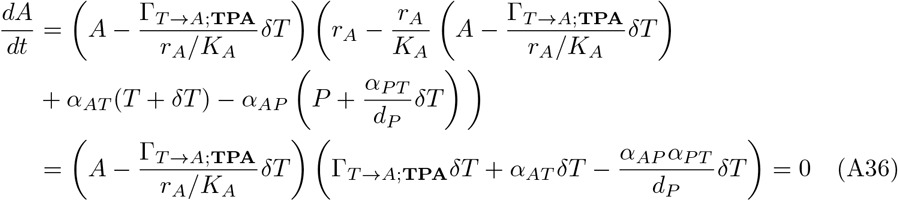

we see that both the PTI and ATI density remain unchanged, so that the system moves away from the tumor-PTI-ATI coexistence steady state. The stability of the steady state when Γ_*T →T* ;**TPA**_ *>* 0 can be checked for each specific case by examining the eigenvalues of the Jacobian matrix, as before.

### A.6 Tumor-PTI-ATI coexistence (TPA) with PTI to ATI conversion (*ω ≥* 0)

We now examine the full steady-state behavior of the model as the ATI to PTI conversion rate is varied. Rearranging the nullclines (setting each of Eqs. 1-3 to zero) for steady states in S_**TPA**_, we find the system of equations

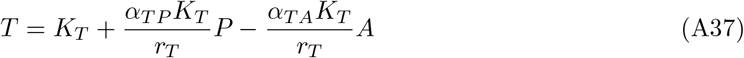

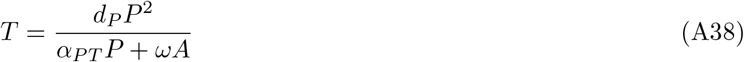

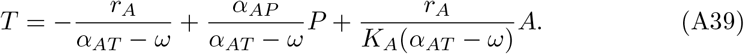

In order to find the steady state, we find the intersection of the two planes (Eqs. A37 & A39) in terms of *P*

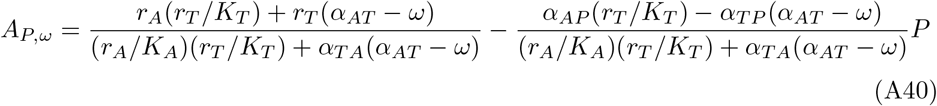

and then look for intersections with Eq. A38 using Eqs. A37 & A40

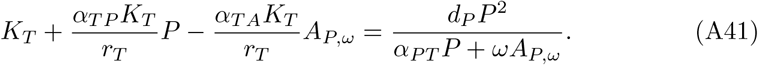

Rearranging Eq. A41 into the form

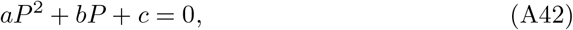

we can easily solve for *P*, and subsequently find *T* and *A*. The bifurcation diagrams in Fig. 2 illustrate the behavior of all the steady states (except for the tumor-only steady state) as *ω* is varied, for parameters where all the steady states of interest are feasible. When *r*_*T*_ − *α*_*T A*_*K*_*A*_ = 0 (first row of Fig. 2), the tumor density is zero at steady state at *ω* = 0. For all *ω >* 0, the system is bistable, with the tumor density either at zero, or for the tumor-PTI-ATI coexistence state, at a positive number. The steady state with non-zero tumor density continuously increases until *ω* reaches 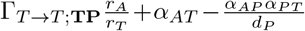 (Eq. A24), when the tumor-PTI-ATI coexistence solution and the tumor-PTI coexistence solution collide, resulting in a transcritical bifurcation (note that we only depict feasible states in Fig. 2). As the rate of tumor death due to ATI cells (*α*_*T A*_) increases, we can see that a saddle node bifurcation gives rise to a stable and an unstable tumor-PTI coexistence solution at a positive *ω*. The *ω* value where this saddle node bifurcation occurs increases with *α*_*T A*_, meaning that tumors need to convert ATI cells to PTI cells at a higher rate in order for a stable tumor population to exist as ATI cells kill tumor cells more rapidly.

When *α*_*T A*_ is large enough, the value of *ω* for which the saddle node bifurcation occurs reaches the value of *ω* for which the tumor-PTI-ATI and tumor-PTI solution collide (Fig. 3). When these two values of *ω* coincide, there is no longer a saddle node bifurcation creating the two tumor-PTI-ATI solutions, but rather a subcritical pitchfork bifurcation giving rise to two unstable tumor-PTI-ATI solutions. At higher *α*_*T A*_ values, the saddle node bifurcation reappears, but now the branch with lower tumor density has a stable region, which is infeasible.

## Appendix B Alternate models: modifications to PTI dynamics

There are several reasonable changes one could make to the model presented in the main text,

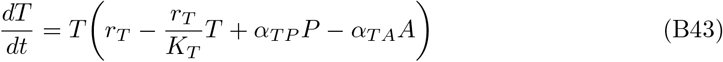

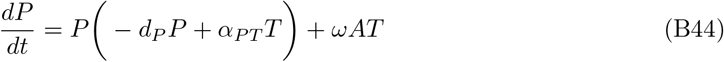

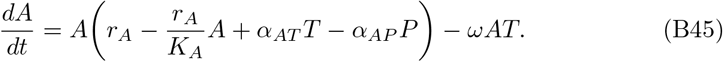

The alternate models we consider involve changes to the PTI dynamics, in this Appendix, and to the ATI dynamics, in Appendix C.

### B.1 Linear PTI death rate

For the changes to the PTI dynamics, we first consider the consequences of changing the PTI death rate to be linear

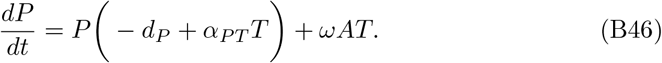

For this model, there can be **T, A, TA**, and **TP** steady states and up to two **TPA** steady states. The actual **T, A**, and **TA** state vectors are the same as in the original model. The stability of **T** can switch as *ω* is changed, due to a transcritical bifurcation resulting from a collision with a stable **TPA** state. The conditions for **T** stability are

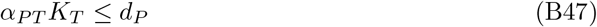

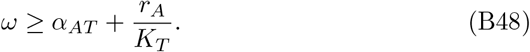

The **A** steady state has the exact same stability conditions as in the original model, while **TA**, which is equal to **TA** in the original model and can still only exist when *ω* = 0, can now be stable if 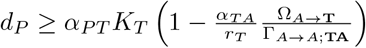. The **TP** state can exist in this alternate model with densities

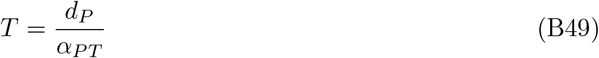

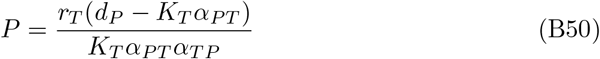

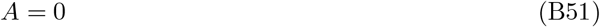

and stability conditions

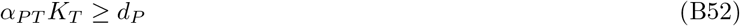

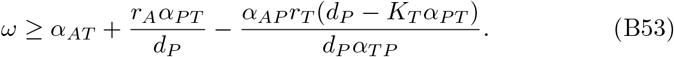

Note that the PTI density of **TP** can only be non-negative when the first stability condition is non-positive. Thus, when **TP** is feasible, it is unstable.

To find the **TPA** states, we can use the approach described for the original model, substituting

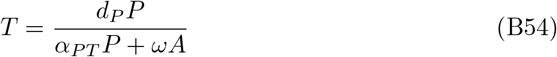

for Eq. A38. This again leads to a quadratic equation in *P*, from which we can solve for two **TPA** steady states.

### B.2 No ATI-independent effect of tumor density of PTI dynamics

Next, we analyze an alternate model where the PTI death rate is nonlinear, but there is ATI-independent effect of tumor density of PTI dynamics (*α*_*P T*_ = 0)

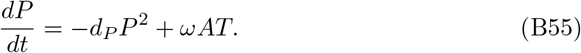

Here, there can be **T, A**, and **TA** steady states and up to two **TPA** steady states. The actual **T, A**, and **TA** state vectors are again the same as in the original model. The **T** state has the single stability condition

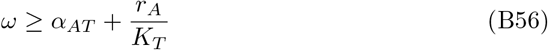

while the **A** state has the same stability criteria as in the previous section. Again, for the **TPA** states, we can use the approach described for the original model, substituting

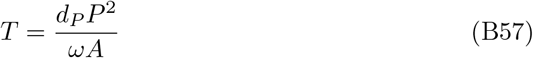

for Eq. A38, yielding a quadratic equation in *P* and up to two **TPA** steady states.

### B.3 Linear PTI death rate and no ATI-independent effect of tumor density of PTI dynamics

Finally, we can make both changes to the PTI dynamics simultaneously, yielding

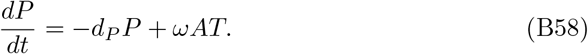

This model yields similar results as in Appendix B.2, except that to find the **TPA** states, we substitute

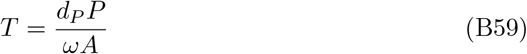

for Eq. A38 and solve the quadratic equation for *P*, yielding up to two **TPA** steady states.

## Appendix C Alternate models: modifications to ATI dynamics

The final modification to our original model that we examine is to the ATI dynamics, where now ATI cells are recruited at a constant rate and die at a constant rate, yielding

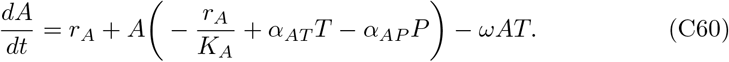

There are now no **T** and **TP** states, because of ATI recruitment (*r*_*A*_). The **A** state and its stability condition are exactly the same as in all other models.

Looking at the nullclines of the model, with each of the four possible models for PTI dynamics, we have

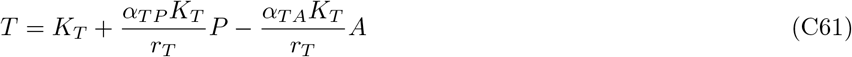

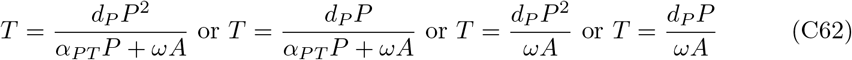

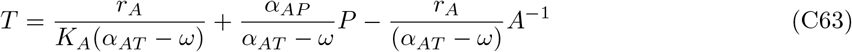

from which we can solve for *P* in terms of *A*

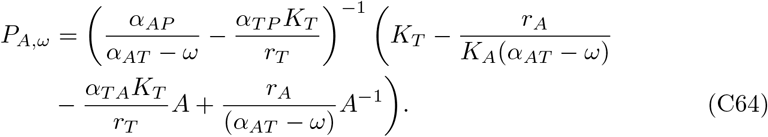

Choosing, for example, the original PTI model dynamics, we end up with the equation

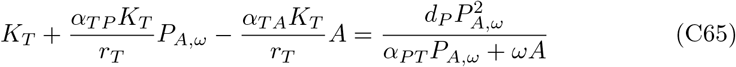

This leads to a degree four polynomial, with at most four roots. The solutions for *A* can then be used to find the **TPA** states.

